# MethylQUEEN: A Methylation Encoded DNA Foundation Model

**DOI:** 10.1101/2024.12.26.630389

**Authors:** Mingyang Li, Ruichu Gu, Shiyu Fan, Yu Fan, Bo He, Jinmin Yang, Yuting Chen, Mengling Xin, Han Wen, Chengqi Yi

## Abstract

DNA 5-methylcytosine (5mC) modification plays a pivotal role in many biological processes, yet 5mC information and pattern hidden behind remains to be explored. Here, we develop **<u>Methyl</u>**ation Language Model based on **<u>Qu</u>**intupl**<u>e</u>** Bidir**<u>e</u>**ctional Tra**<u>n</u>**sformer (MethylQUEEN), a novel pre-trained DNA methylation foundation model capable of sensing methylation states and covering the genome-wide methylation landscape. Through tailored methylation-prone pre-training, MethylQUEEN effectively captured epigenetics information hidden within the DNA sequences: it accurately traces DNA’s tissue-of-origin, and successfully recovers the expression profile through methylation states. Integrative analysis on MethylQUEEN’s attention scores also enables us to reveal the unique methylation status of a tissue for precise disease detection, and identifying key regulatory 5mC sites for disease intervention. As a result, MethylQUEEN signifies a new paradigm in methylation analysis for various biological problems. Besides, our study demonstrates the effectiveness of directly integrating methylation information into pre-training, offering new perspectives and methodologies for a range of methylation-related biological processes. It serves as an initial exploration for the development of more comprehensive epigenomic models.

## Introduction

DNA 5-methylcytosine (5mC) methylation is the most common epigenetic modification that involves the addition of a methyl group to the fifth position of cytosine residues, primarily within CpG dinucleotides in mammalian genomes^1^. 5mC is pivotal in regulating gene expression, genomic stability, and cellular differentiation by affecting chromatin structure and transcriptional activities^2^, especially in development and disease^3,4^. Aberrant methylation patterns, such as hypermethylation of tumor suppressor genes or global hypomethylation, are frequently associated with various cancers and other diseases^4–6^. Advances in high-throughput sequencing technologies have facilitated the characterization of 5mC at single-base resolution, enriched our understanding of its dynamic role in cellular function and epigenetic inheritance^7^.

Among the sequencing technologies for detecting 5mC signals, BS-seq has been recognized as the gold standard, which generated high-quality datasets for constructing comprehensive DNA methylation maps and uncovering their underlying mechanisms. However, unlocking the hidden insights within these datasets and efficiently extracting the epigenetic information from vast sequence spaces remain significant challenges, calling for innovative advancements in computational methods. Recently, large-scale pre-trained models have exhibited remarkable capability in natural language processing^8,9^. Similarly, nucleic acid sequences, as the primary carriers of genetic information, are well-suited to be processed as ‘biological languages’ of nature. Numerous outstanding models have been developed for nucleic acid sequences, including Evo^10^ and DNABERT^11^ for DNA Uni-RNA^12^ and RiNALMo^13^ for RNA. Existing research either included epigenetic information only during fine-tuning stage^14^ or directly into pre-training but in the form of reduction to beta values^15,16^, which ignores the vast information hidden in a single biological sample. There is an urgent need to fill this gap towards a foundation model that directly incorporates epigenetic information from BS-seq data during the pre-training phase.

In this study, we developed the **<u>Methyl</u>**ation Language Model based on **<u>Qu</u>**intupl**<u>e</u>** Bidir**<u>e</u>**ctional Tra**<u>n</u>**sformer (MethylQUEEN), a methylation-state-aware pre-trained model based on high-quality genome-wide 5mC methylation data. This model promotes a scalable and easy-to-expand modification encoding scheme. Aimed at capturing sequence features and their hidden epigenetic information, MethylQUEEN was pre-trained on over 500 million high-quality assembled BS-seq sequences from various human tissues and with a specially calibrated framework^17^ to achieve efficient sequence representation. As a result, MethylQUEEN demonstrated superior performance in a wide range of methylation-related downstream tasks (**Figure 1a**). As the first foundation model directly trained on BS-seq data, MethylQUEEN signifies a paradigm shift in methylation analysis, providing researchers with a more powerful and efficient tool to analyze methylation data and uncover the underlying biological mechanisms based on the comprehensive knowledge encoded in the model’s embeddings and attention scores, paved way for early disease detection and precise medicine.

**Figure 1.**
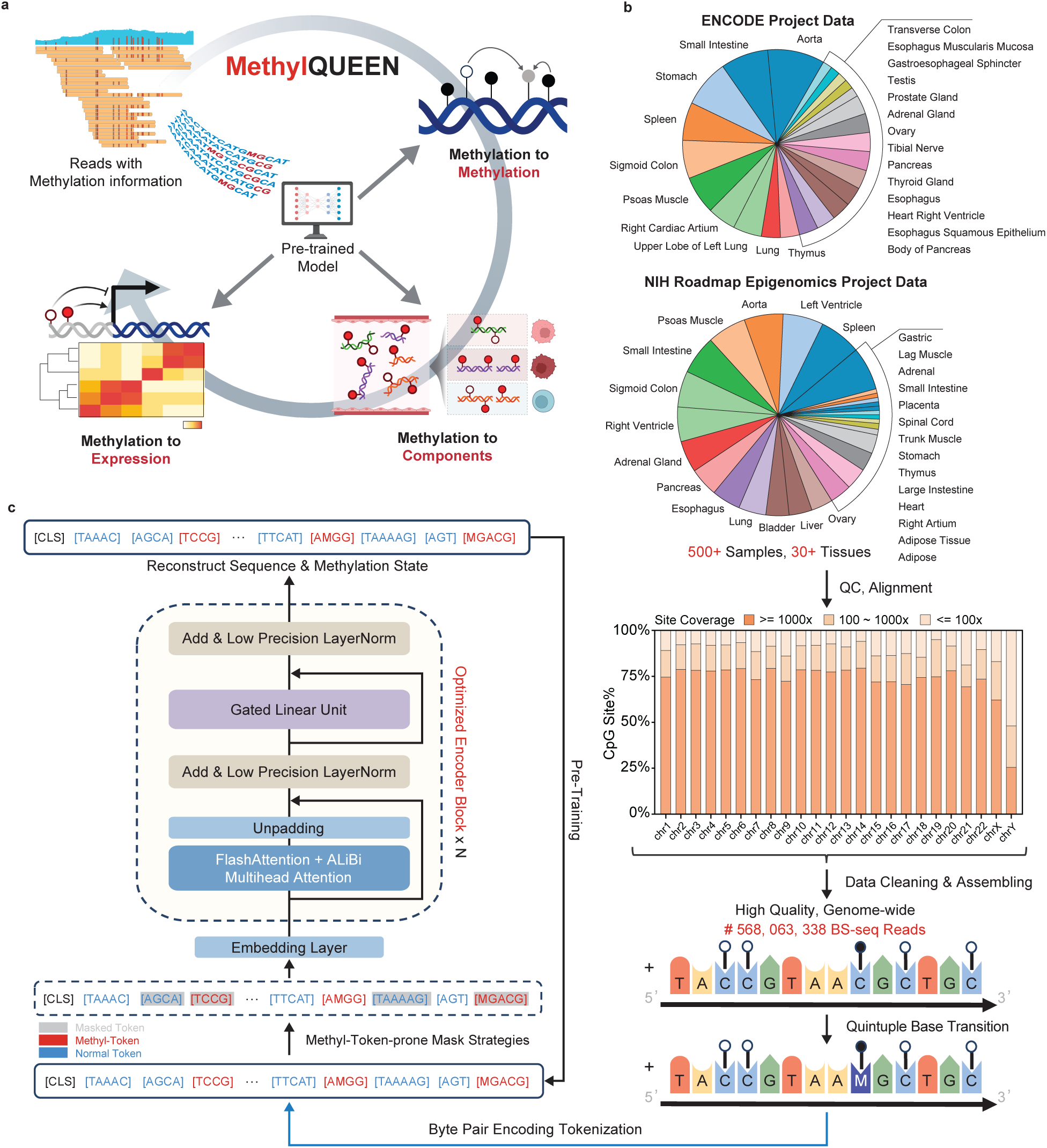
Introduction of MethylQUEEN, and its various pretraining data. **(a)** MethylQUEEN takes methylation sequencing data as input and generates high-dimensional representations that integrate sequence and methylation features. After fine-tuning, it enables accurate predictions for methylation-related biological processes, such as gene expression regulation, tissue classification, and disease monitoring. **(b)** The pretraining dataset for MethylQUEEN was derived from two renowned DNA sequencing projects, ENCODE and NIH Roadmap Epigenomics Mapping program, incorporating BS-seq results from over 500 samples spanning more than 30 tissues. This dataset provides coverage of most CpG sites with over 1,000 sequencing reads per site. After data cleaning based on sequence length and quality, more than 500 million high-quality DNA sequences containing methylation information were retained. **(c)** MethylQUEEN was trained using a methylation-specific Byte Pair Encoding(BPE) tokenizer designed for the Quintuple genome to encode DNA sequences with methylation information. The model adopts the Mosaic BERT framework, which integrates advanced strategies such as FlashAttention and ALiBi positional encoding and specifically adapted for DNA methylation scenarios and employs a BERT-like training scheme to ensure efficient and accurate reconstruction of both sequences and methylation information.

## Results

### MethylQUEEN leverages advanced pre-training strategies for efficient methylation representation

To ensure comprehensive coverage of the DNA methylation space, we gathered over 500 biological samples from two prominent DNA sequencing initiatives: the ENCODE^18^ and the NIH Roadmap Epigenomics^19^. After conducting quality control and sequence alignment, we achieved deep coverage of all CpG sites across the whole human genome. This process yielded 568,063,338 high-quality methylation-encoded sequences, which were cleaned, assembled, and transformed into a quintuple format to encode methylation states, with methylated cytosine presented as letter ‘M’. These formatted sequences were then tokenized using a Byte Pair Encoding (BPE) tokenizer as input for the model (**Figure 1b**).

To optimize model performance and facilitate downstream analyses, we define all possibly methylation related patterns, such as “CG”, methylated base “M”, and the token “C” (accounting for unconventional methylation patterns) as methyl-tokens. The MosaicBERT^17^ module served as the foundational framework, with a methyl-token-prone pre-training strategy designed to enable the model to simultaneously recover sequence information and methylation states from context (see Methods). Additionally, advanced techniques such as Attention with Linear Biases (ALiBi)^20^ and flash attention^21^ were integrated to maximize training efficiency and enhance the model’s representational capacity (**Figure 1c**).

After pretraining, all methyl-tokens were clustered into a distinct group (**Figure 2b**) and the immune-tissue-specific methylation information can be directly captured from the massive sequencing data (**Figure 2d**). This demonstrates MethylQUEEN’s ability to learn hidden methylation patterns embedded in the sequences. We next utilized MethylQUEEN to perform methylation state imputation, aiming to reduce the uncertainty in methylation state annotations (**Figure 2e**). The results showed that, even with datasets of varying scales, MethylQUEEN could effectively reconstruct methylation states with high accuracy (**Figure 2f**).

**Figure 2.**
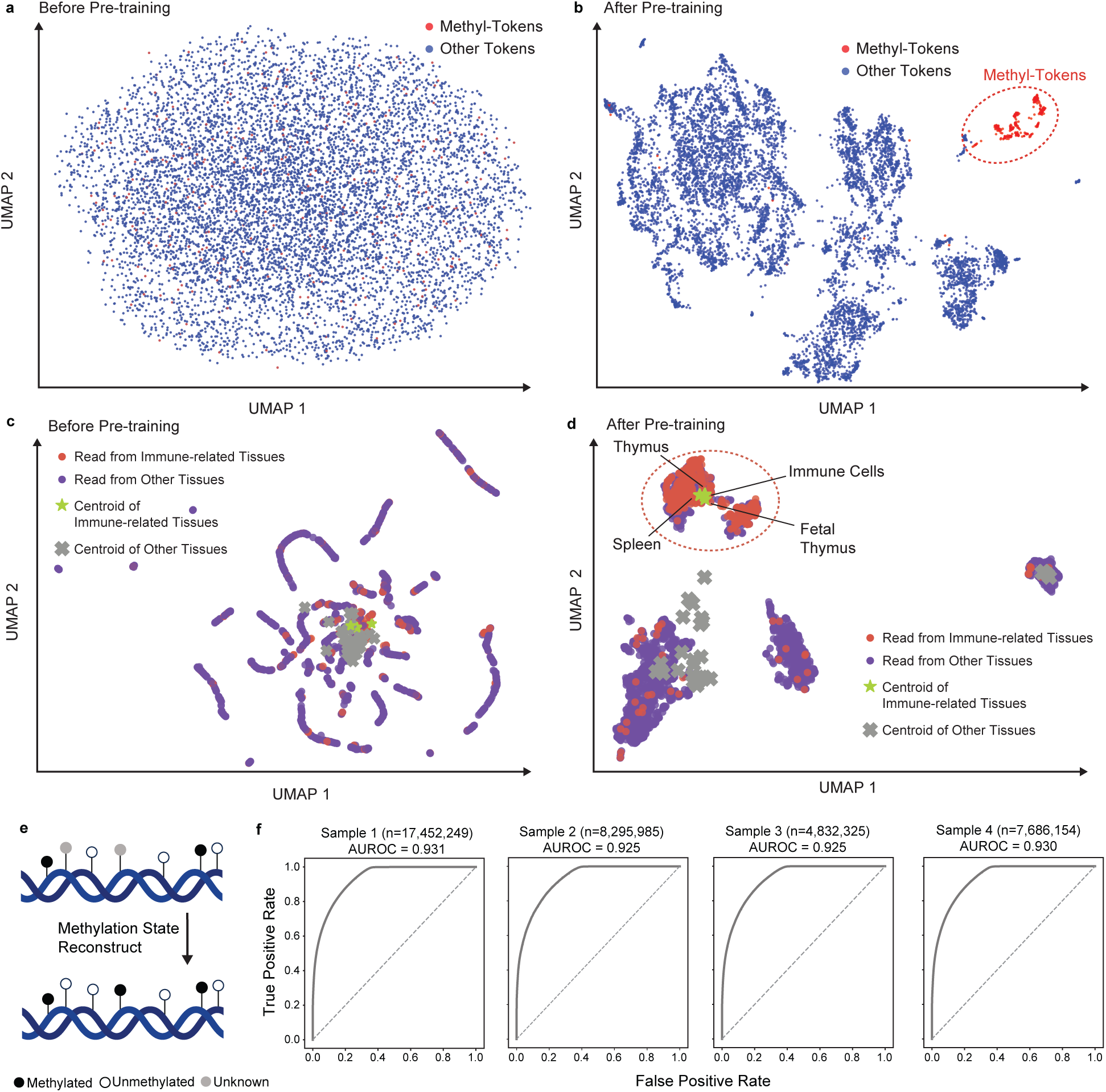
Pre-trained MethylQUEEN achieved zero-shot methylation analysis. **(a, b)** UMAP projection of MethylQUEEN embeddings for all tokens in the BPE tokenizer before and after pre-training. **(c, d)** UMAP projection of MethylQUEEN embeddings for reads from DMR across different tissues before and after pre-training. **(e)** Reconstruction of ambiguous methylation data using MethylQUEEN to improve accuracy for imprecisely sequenced regions. **(f)** Fine-tuned MethylQUEEN’s receiver operating characteristic(ROC) curve and confusion matrix to evaluate the methylation state reconstruction performance on various samples(The methylation sequencing data were obtained using PacBio HiFi CCS, with methylation states determined via electrical signals).

### MethylQUEEN achieved effective sequence classification in various scenarios

Past research on DNA methylation predominantly employed a methodology that compares methylation differences between two components to identify differentially methylated sites (DMSs) or regions (DMRs). These components often include cancerous versus healthy tissues in cancer research or senescent versus proliferating cells in cellular senescence research. Broadly, these comparisons can be categorized as treatment components versus control components. The primary biological objectives typically involve identifying key methylation markers associated with abnormal gene expression, disease detection, and other related phenotypes. However, many existing algorithms reduce BS-seq reads to beta values, losing critical read-level information, prompting an increasing focus on the read-level analysis of identified DMRs, especially in liquid biopsy. We defined a method that directly assigns methylation-encoded reads to the components of interest, referred to as methylation-to-components (M2C). This read-level analysis enables more accurate estimations of component fractions in mixed environments, such as plasma, and facilitates improved detection of cancers or other diseases through liquid biopsy and bulk tissue deconvolution (**Figure 3a**).

**Figure 3.**
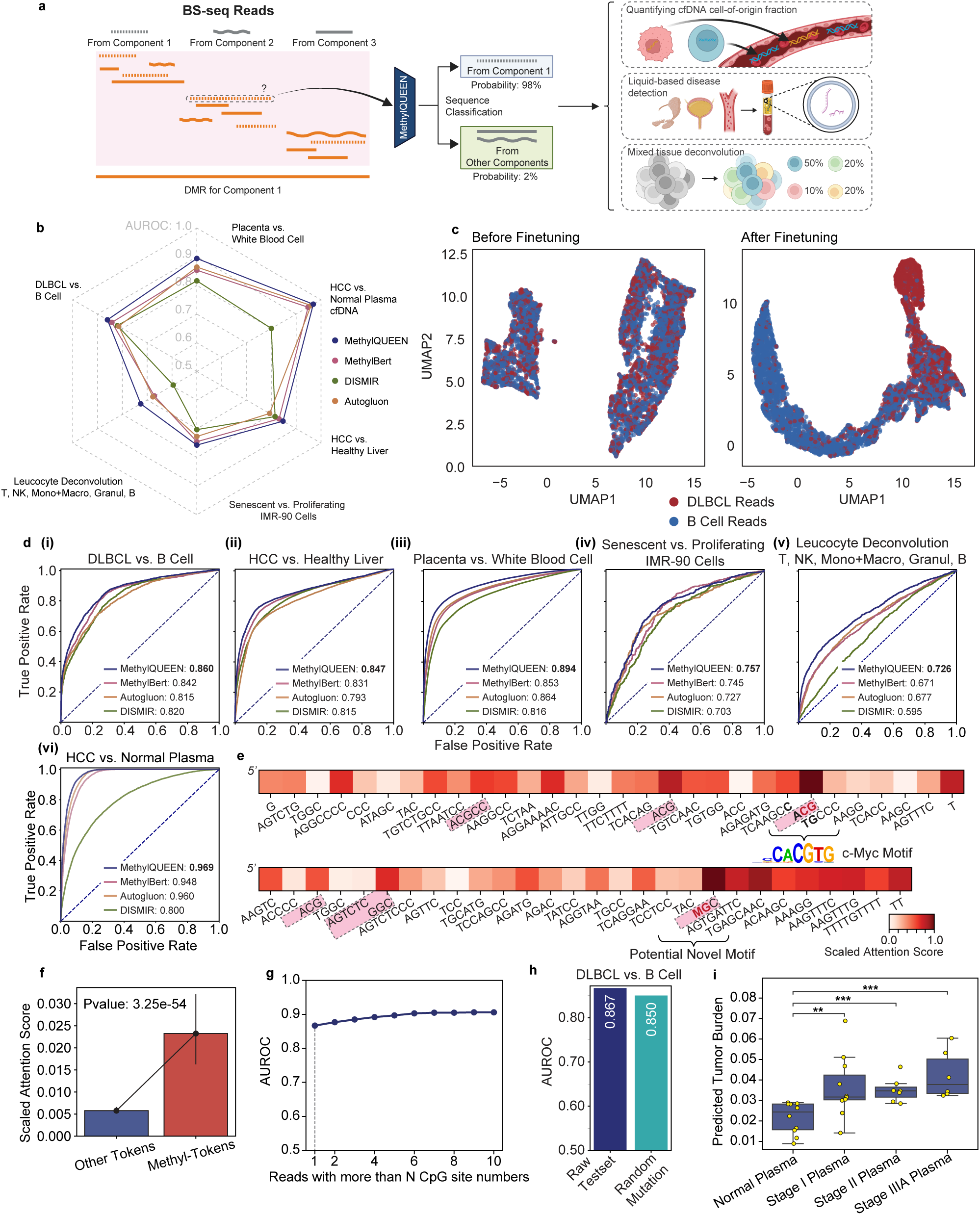
MethylQUEEN improves the precision of methylation sequence annotation. **(a)** Schematic representation of the Methylation-to-Component (M2C) task. Within component-specific DMRs, the origin of reads can be identified using fine-tuned MethylQUEEN. M2C tasks can be applied in biological and clinical scenarios such as estimating cfDNA composition in body fluids, disease detection, and deconvolution of mixed tissues or cell types. **(b)** Radar plot showing the area under ROC curves (AUROC) for fine-tuned MethylQUEEN compared to other previously reported models on test sets across six M2C tasks: Hepatocellular carcinoma (HCC) vs. Healthy liver tissue or normal plasma, Placenta vs. White blood cells, Diffuse large B-cell lymphoma (DLBCL) vs. Normal B cells, Senescent IMR-90 cells vs. Proliferating IMR-90 cells, and Leukocyte deconvolution (T cells, NK cells, Monocytes + Macrophages, Granulocytes, and B cells). **(c)** UMAP projection of reads from placenta and white blood cell components, visualized before and after fine-tuning with MethylQUEEN. **(d)** ROC curves of fine-tuned models on test sets for four M2C tasks. **(e)** Heatmap showing the scaled attention scores of two reads. **(f)** Scaled attention scores for two types of tokens. **(g)** Line chart illustrating the relationship between the minimum number of methylation sites (CpG sites) in reads and AUROC values. **(h)** AUROC values of fine-tuned MethylQUEEN on the original test set and a randomly mutated test set for the DLBCL vs. Normal B cells task. **(i)** Estimation of HCC tumor burden in clinical plasma samples from HCC patients at different stages and healthy controls. **p < 0.01, ***p < 0.001.

In this study, we comprehensively evaluated MethylQUEEN’s performance in M2C tasks and achieved the best performance across various downstream tasks in diverse scenarios (**Figure 3b**). In detail, we took the difference between reads from DLBCL and B cells as an example to demonstrate that fine-tuning effectively separated reads from two components, as evidenced by their clear division after fine-tuning (**Figure 3c**). Furthermore, in other important biological tasks (see detail in Methods), MethylQUEEN consistently outperformed other models. It achieved up to a 6.03% increase in the area under the ROC curve (AUC) (**Figure 3d**). These findings highlight MethylQUEEN’s efficiency and superiority in M2C tasks.

We conducted an in-depth analysis of the attention scores to further explore hidden insight behind the optimal performances. For instance, in the DLBCL vs. normal B cells task, the model captured a methylation-encoded sequence located in the promoter region of the *MSX1* gene (**Figure 3e, top**). *MSX1* is known to play a role in early B-cell development and the malignant transformation of DLBCL^22^. Within this sequence, a CpG site in the context of CA<u>CG</u>TG displayed the highest attention score, namely the model believes it’s the most relevant site. Meanwhile, this CpG context matched the motif of c-Myc, suggesting that this site may be critically involved in DLBCL development by influencing the expression of the *MSX1* gene through recognition by the c-Myc transcription factor (TF) motif. Another sequence within the *FOXQ1* gene, a key player in tumor pathogenesis^23^, showed a similar high attention score at the CpG site located in a sequence context (AC<u>CG</u>CA) that does not match any known TF binding motifs (validated in HOMER and Tomtom databases), potentially representing a novel motif (**Figure 3e, bottom**).

Furthermore, we performed a statistical analysis of token attention scores across the entire test set and observed that methyl-tokens received significantly higher attention than other tokens (**Figure 3f**). This finding clearly indicates MethylQUEEN’s sensitivity to methylation information when analyzing methylation-encoded sequences. Additionally, we investigated the relationship between classification efficiency and the number of CpG sites in a sequence. We found that the presence of a single methylation site in the sequence is sufficient for MethylQUEEN to achieve effective classification, and an increased number of CpG sites per read only mildly enhances classification performance (**Figure 3g**). Based on this observation, we suggest that during dataset construction in these scenarios, filtering sequences based on the presence of at least one CpG site is sufficient.

To assess MethylQUEEN’s robustness, we introduced random mutations to non-CpG sites in the test dataset reads. Following these mutations, the AUROC showed only a slight decrease, demonstrating its robustness in M2C tasks (**Figure 3h**).

Finally, to demonstrate the potential of MethylQUEEN for liquid biopsy applications in complex clinical cancer detection after validating the high performance of MethylQUEEN, we analyzed a clinical dataset containing targeted BS-seq libraries from normal plasma cfDNA and plasma cfDNA from patients with HCC at various stages. The predicted tumor burden for stage I HCC, an early stage that is notoriously difficult to detect through liquid biopsy^7^, showed a significant increase compared to normal plasma samples (**Figure 3i**). These findings highlight the capability of MethylQUEEN to detect cancers at extremely early stages, suggesting its potential utility in future clinical practice for early cancer diagnosis.

### MethylQUEEN accurately predicts gene expression levels and identifies key regulatory CpG sites

Since the discovery of DNA methylation, its significant regulatory role in gene expression has remained a crucial area of research, driving studies across numerous biological and clinical domains. These include gene silencing and activation mechanisms^2^, cellular differentiation and cell fate decisions^24^, malignant tumor proliferation^6^, individual aging^25^, and so on. DNA methylation largely determines various biological phenotypes through its regulation of gene expression and the consequent gradual transmission of regulatory information along the central dogma.

Following this rationale, we attempted to predict gene expression *in silico* via methylation information (methylation-to-expression, M2E) based on MethylQUEEN. Such computational prediction models could facilitate integrative analyses of large-scale omics datasets, paving the way for advances in personalized medicine, such as using epigenomic editing technologies^26,27^ and in system biology research (**Figure 4a**).

**Figure 4.**
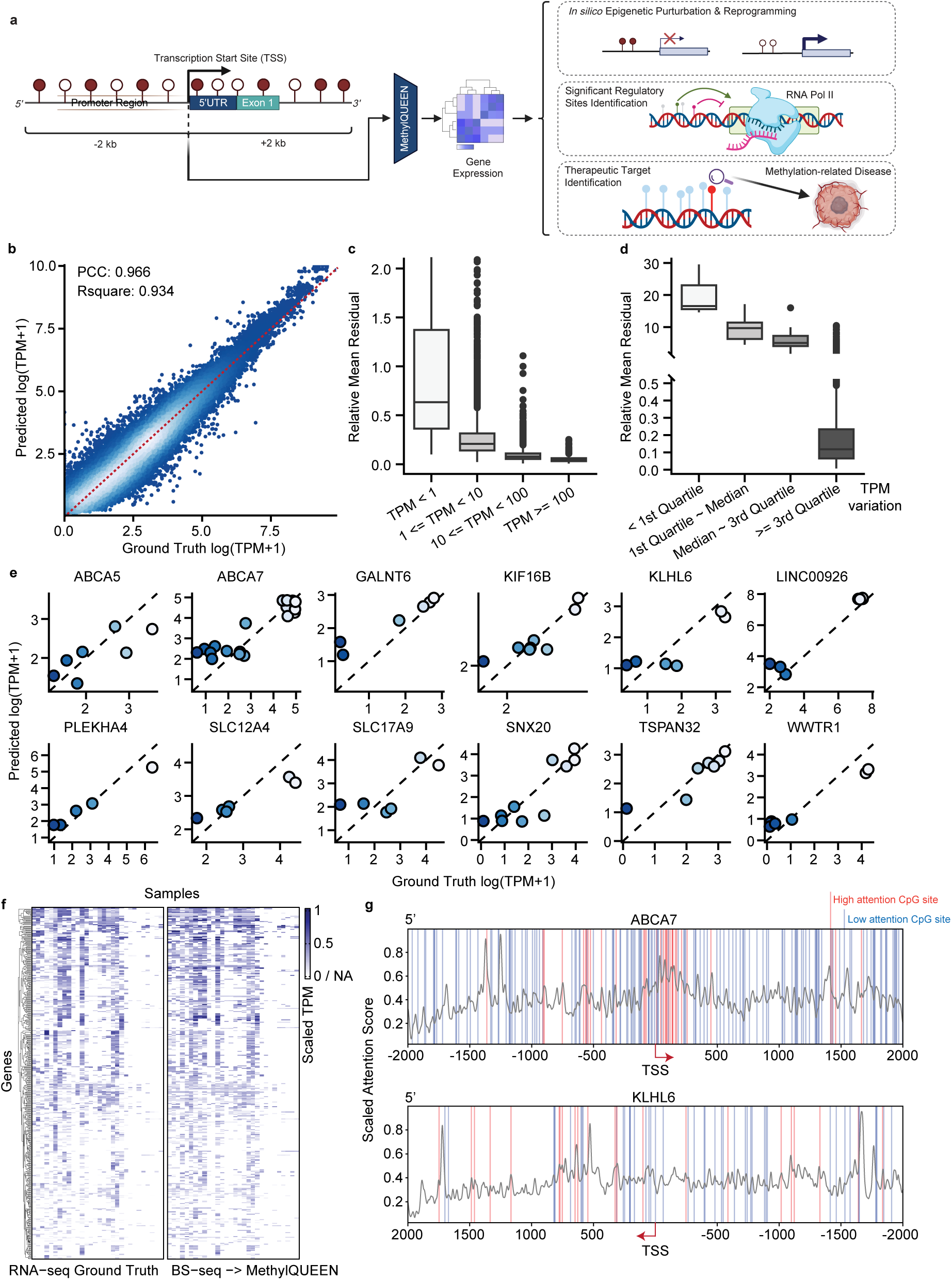
MethylQUEEN uncovers methylation regulation on expression pattern. **(a)** Schematic representation of the Methylation-to-Expression (M2E) task. Methylation status within ±2 kb of the transcription start site (TSS) is input into the fine-tuned MethylQUEEN, which predicts an estimated expression value based on the current methylation state. M2E tasks can be applied to in silico methylation perturbation or reprogramming, identification of significant regulatory CpG sites, and therapeutic target CpG sites for DNA methylation editing. **(b)** Scatter plot comparing ground-truth expression values from RNA-seq with predicted expression values derived by MethylQUEEN from BS-seq data. **(c)** Relative mean residuals at different mean expression TPM values. **(d)** Relative mean residuals at different quantiles of gene expression variance. **(e)** Comparison of ground-truth expression values and predicted expression values for selected example genes. **(f)** Heatmap comparing gene expression profiles obtained from RNA-seq with reconstructed gene expression profiles predicted by MethylQUEEN based on BS-seq data. **(g)** Scaled attention scores of MethylQUEEN for ±2 kb TSS-flanking regions of the ABCA7 and KLHL6 genes. The plot highlights CpG sites with high scaled attention scores (red bars, score ≥ 0.5) and low scaled attention scores (blue bars, score < 0.5). Grey lines show the smoothed attention score distribution across the region.

We utilized public datasets comprising matched BS-seq and RNA-seq data from the same biological samples to fine-tune MethylQUEEN for the M2E task. The model was trained to predict gene expression levels (TPM) using methylation-encoded sequences located within ±2 kb around the transcription start site (TSS) of genes (**Figure 4a**). The region ±2 kb around the TSS involves genomic elements like promoter, 5’UTR and first exon, where the promoter is the most influential element where methylation regulates^2^. On the test set, the predicted TPM values strongly correlated with RNA-seq-detected TPM values, achieving a Pearson correlation coefficient (PCC) of 0.966 and an R-squared value of 0.934 (**Figure 4b**).

Further, we revealed that genes with higher mean expression values and greater expression variation were easier to predict (**Figure 4c, d**). Specifically, genes with expression value variations exceeding the third quartile were significantly easier to predict than others. These findings highlight the necessity of incorporating more diverse training datasets collected from various tissues and conditions in future research.

To enhance the diversity of the training dataset, we integrated methylation data from the ENCODE and NIH Roadmap Epigenomics Project datasets, which include samples from a wide range of tissues. To obtain paired gene expression values for training, we downloaded the Genotype-Tissue Expression (GTEx) dataset^28^, which provides tissue-specific gene expression data. Using this dataset, we constructed a normal distribution of gene expression values for each tissue (see Methods) and sampled matched expression values for each methylation-encoded sequence. After expanding the training dataset, we observed that MethylQUEEN achieved high prediction accuracy for genes associated with DNA methylation or other biological significances (**Figure 4e**). By comparing gene expression profiles derived from RNA-seq and MethylQUEEN’s predictions based on BS-seq data, we found a high degree of similarity between the predicted profiles and those obtained via RNA-seq (**Figure 4f**). These results demonstrate MethylQUEEN’s remarkable ability to predict gene expression profiles based on methylation data accurately. This underscores its potential for future applications in *in silico* methylation perturbation studies and investigations of methylation-mediated gene regulation.

Finally, we analyzed the attention score profiles of the *ABCA7* and *KLHL6* gene regions during MethylQUEEN analysis. The *ABCA7* gene is strongly associated with Alzheimer’s disease. It is implicated in DNA methylation regulation^29^, while *KLHL6* has been reported to play a role in cancer development and is also related to DNA methylation^30^. When MethylQUEEN predicted the expression values based on the methylation information of these two genes, we observed that certain CpG sites received significantly higher attention scores while others did not (**Figure 4g**). The methylation states of CpG sites with higher attention scores had a greater influence on expression predictions, suggesting that these CpG sites have stronger regulatory potential. This indicates that MethylQUEEN not only accurately predicts expression values but also identifies CpG sites that are likely critical for proximal methylation regulation. Such CpG sites could serve as potential therapeutic targets in the future.

## Discussion

In this study, we have developed MethylQUEEN, a large-scale pre-training model for DNA sequences with methylation information. It has been trained on over 500 million high-quality assembled BS-seq sequences and demonstrated superior performance in various downstream tasks related to methylation state. Supported by extensive high-quality methylation data, MethylQUEEN has the potential to transform our understanding of methylation from isolated observations to interconnected insights, from local contexts to a holistic perspective. More importantly, we systematically demonstrated that directly incorporating epigenomic information into the pre-training process yields better performance compared to pre-training solely on genome sequences. We also provide the easy-to-expand modification encoding method for subsequently integrating diversity modifications in the future pre-training, such as m6A in RNA, which could provide a theoretical foundation for incorporating more epigenetic data in future studies and significantly broaden the scope of data that can be utilized for pre-training.

In summary, MethylQUEEN provides a high-performance and efficient framework with the integration of an expanded nucleotide genome encoding epigenetic information. MethylQUEEN’s embeddings and attention scores offer a novel methodological framework for methylation-related analyses, paving the way for a deeper exploration of the mechanisms by which DNA methylation regulates biological processes. Our method for encoding modifications to nucleotide sequences is simpler and easier-to-expand than adding extra embedding vectors^14–16^ (named the extra embedding vector solution), as it can be readily extended to accommodate various types of nucleotide modifications in the future by simply expanding the tokenizer vocabulary. Furthermore, integrating multiple computational optimizations enables efficient training and fine-tuning of MethylQUEEN. These advancements lay a solid foundation for developing comprehensive epigenomic models encompassing diverse types of modifications.

In the future, we plan to optimize MethylQUEEN from both data and model perspectives, further exploring the application of data-driven pre-trained models in epigenetics. On the data front, we aim to expand the pre-training data space by incorporating various modifications. Encoding multiple modification types could help us to unveil intra-modification direct or indirect crosstalk (like DNA 5mC and RNA m6A), conduct in-depth analyses of biological problems involving multiple modification types, identify key modification types in a mechanism, imputing multi-modification patterns, etc. MethylQUEEN’s efficient representations will enable the alignment and fusion of these diverse data modalities. On the model front, generative models have already achieved significant success in the biological domain^31–33^. Generative models in the methylation field can also play an important role and significance in more challenging issues such as methylation pattern research and methylation function annotation. Balancing the trade-off between representational power and generative capacity will be a key focus for architectural optimization in the next stages of MethylQUEEN’s development.

## Methods

### Pre-training dataset collection, analysis and cleaning

The pre-training dataset included whole-genome bisulfite sequencing (WGBS) samples from the ENCODE Project^18^ and the NIH Roadmap Epigenomics Project^19^. All available tissue data from *Homo sapiens* were downloaded and processed using a standard BS-seq alignment pipeline available on GitHub (https://github.com/Wubeizhongxinghua/DNA_5mC_analysis_pipeline). Briefly, the pipeline first performed quality control using trim_galore (v0.6.10) with the following parameters: -j 20 -q 30 --phred 33 --length 35 --stringency 3. Filtered reads were then aligned to the GRCh38 human genome using bwameth (v0.2.7) with default parameters. The aligned reads were filtered through the following steps: 1) Duplicate alignments were removed using picard.jar MarkDuplicates (v2.27.5) with default parameters. 2) For paired-end reads, only those with flags 99, 147, 83, or 163, and for single-end reads, only those with flags 0 or 16, and with a MAPQ score ≥ 60 were retained. 3) Reads with more than three CH mismatches (for flags 99, 147, and 0) or three GD mismatches (for flags 83, 163, and 16) were considered incompletely converted by bisulfite and were removed. The remaining reads constituted the final processed dataset. In these reads, unmethylated cytosines (C) were labeled as “C” (converted to T), while methylated cytosines were labeled as “M” (unaltered Cs). The processed BS-seq reads had lengths ranging from 98 to 151 bp, which were unsuitable for pre-training. To address this, short reads were assembled into longer reads (mean length > 600 bp) by overlapping matched methylation states within each biology sample. After assembly, reads containing more than 32 Ns or exceeding 4000 bp in length were discarded. Finally, high-quality assembled reads from the forward strand (flags: 99, 147, 0) were selected as the assembled read pool. These reads were used for byte-pair encoding tokenizer training and MethylQUEEN pre-training.

### MethylQUEEN architecture and pre-training

MethylQUEEN adopted the MosaicBERT architecture^17^, with specific modifications designed to accommodate the unique characteristics of methylation data. These adjustments provided significant improvements in computational efficiency and memory usage compared to the original BERT architecture. Specifically, MethylQUEEN incorporated the following features: 1) Positional embeddings were replaced with Attention with Linear Biases (ALiBi)^20^, enabling better performance on sequences with a wide range of lengths. 2) FlashAttention^21^ and Low Precision Layer Normalization were implemented to optimize computational efficiency and reduce memory usage. 3) The ReLU activation function was replaced with the Gated Linear Unit with GeLU (GeGLU)^34^, which has been shown to improve the performance of Transformer models. 4) Low-Rank Adaptation (LoRA)^35^ was incorporated during fine-tuning to achieve parameter-efficient training.

MethylQUEEN utilized byte-pair encoding (BPE) for tokenization. To train the tokenizer, 5% of the reads from the assembled read pool were selected, and the HuggingFace tokenizers module was used with a vocabulary size of 8192. The resulting vocabulary prioritized tokens containing CG or M, as well as the single-base token C, which we referred to as methyl-tokens.

Then, we conducted a multi-step pre-training procedure, referred to as the methyl-token-prone pre-training method.

In the first step, standard masked language modeling (MLM) with a masking probability of 0.15 was performed for basic from-scratch pre-training. The training was conducted for 300,000 steps with a batch size of 2048, a peak learning rate of 1e-4, a linear warmup scheduler over 50,000 steps, Adam optimizer with beta2=0.98, and a weight decay of 1e-2.

During the second step, non-methyl-tokens were assigned a masking probability of 0%, while methyl-tokens were assigned a masking probability of approximately 96.98% (1 - total frequency of non-methyl-tokens in the corpus). MethylQUEEN was trained for an additional 280,000 steps during this step with smoothed label during CE loss calculation, using the same hyperparameters as in the first step. All pre-training steps were conducted using 8 NVIDIA A800 80GB PCIe GPUs.

### Methylation state imputation

The PacBio HiFi sequencing libraries dataset was used to conduct the methylation state imputation experiment. For each HiFi sequence, every methyl-token was masked with a probability of 20%. The methylation state of the masked methyl-tokens was then determined by summing the probabilities of methylated methyl-tokens (tokens with “M”) or non-methylated methyl-tokens (tokens with “CG” and the single-base token “C”).

### Calling for differentially methylated regions

To identify tissue-specific differentially methylated regions (DMRs), we employed a one-vs-all strategy, comparing a target tissue against all other tissues to pinpoint tissue-specific DMRs. Methylation information at CpG sites for each sample was initially extracted from BAM files using Methyldackel(https://github.com/dpryan79/MethylDackel) (v0.5.2) with the parameters -q 10 -p 10 --mergeContext. Since some samples had relatively low sequencing depths and the total number of samples was large, we merged tissues with more than three samples to ensure that each tissue retained three representative samples.

Subsequently, Spearman correlation coefficients were calculated for methylation patterns between tissues to assess inter-tissue correlations. Based on these results, tissues with high correlations were further merged. Preliminary DMR calling was then performed using Metilene (v0.2.8) with the parameter --mincpgs 3 to identify raw DMRs.

Building upon these raw DMRs, we applied additional filtering using the one-vs-all strategy. Specifically, we calculated the differences in methylation rates between the target tissue samples and those of each other tissue. Regions where the methylation rate of the target tissue consistently differed from all other tissues by a predefined threshold were retained as tissue-specific DMRs.

### Construction of datasets for M2C task

BS-seq sample libraries were first aligned using the aforementioned pipeline. The reads were then filtered through the following steps: 1) Reads located within DMRs were selected. For each read, regions outside the DMRs were trimmed, and only the portions inside the DMRs were retained. 2) Trimmed reads with a length ≥ 75 bp and flags of 99, 147, or 0 were selected. 3) Reads without CpG sites were removed. 4) Reads were split into training, validation, and test sets based on their respective samples. For datasets without enough samples, the split of training and validation set is split by reads, not samples.

After these steps, some reads with identical sequences but differing component labels or the same label may have existed. Reads with the same labels were merged into one sequence, while those with different labels were removed. Additionally, if any read was shared between the training, validation, and test sets, it was deleted to avoid data leakage.

For robustness testing, reads were randomly selected from the test dataset, and 2 or 3 SNPs or indels were introduced into each read. This process was repeated 250,000 times, ensuring label balancing throughout.

### Methylation to components task benchmarking

For benchmarking, MethylBERT^14^, DISMIR^36^, and AutoGluon^37^ were selected. For MethylQUEEN fine-tuning, Low-Rank Adaptation (LoRA)^35^ was applied using the HuggingFace PEFT module with a learning rate of 1e-5, a linear warmup ratio of 6%, and a dropout rate of 0.1. MethylBERT (v2.0.1) was fine-tuned using the pre-trained model provided on HuggingFace (https://huggingface.co/hanyangii/methylbert_hg19_12l), with hyperparameters provided by authors. For fine-tuning the DISMIR model, we followed the data preprocessing guidelines and model architecture specified by the authors (https://github.com/XWangLabTHU/DISMIR). Specifically, 66 bp-long reads were converted into a one-hot matrix for model input. The model was trained using default hyperparameters, including a learning rate of 0.05, 150 epochs, and a batch size of 128, across the various benchmark datasets. For AutoGluon training, quintuple-base (ACGTM) sequence data were one-hot encoded. The AutoGluon framework (v1.1.0) was then used with default parameters to train machine learning models on the encoded datasets. Validations were performed for all the models per training epoch, and checkpoint with the highest performace on the validation set was used. Inference was then performed on the test sets, and the results were reported. All models were trained using Cross Entropy Loss. Steps requiring GPUs were executed on NVIDIA L40 40GB PCIe GPUs.

### Tumor burden estimating in plasma samples

To estimate the component fraction or purity in genomic regions based on sequencing data, we employed a pipeline involving likelihood maximization, skewness adjustment, and optimization^14^.

Firstly, the tumor purity for all reads within a genomic region (DMRs, in this case) was estimated by maximizing the likelihood of observing the given sequencing probabilities. The likelihood function was defined as:

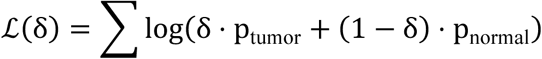

Here δ represents the tumor purity of the region, p_tumor_ is the probability of reads originating from tumor as estimated by MethylQUEEN, and p_normal_ is the probability of reads originating from normal cells. The tumor purity δ was estimated by solving:

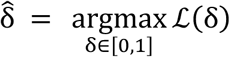

This optimization was performed using the bounded scalar minimization method provided by the scipy.optimize module, specifically using minimize_scalar function.

Secondly, the skewness of the estimated purities across regions was minimized to improve the robustness of the purity estimation. Skewness was calculated using the statistical moment by

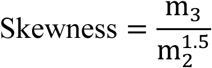

Here, *m*_3_ and *m*_2_ are the third and second moments of the distribution of adjusted purities, respectively. The adjusted purities for each region were defined as:

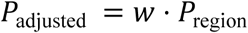

where *w* is a weight vector that scales the estimated region purities P_region_.

Thirdly, the optimal weights *w* were determined by minimizing the skewness of the adjusted purities using the L-BFGS-B method. The optimization problem was formulated as:

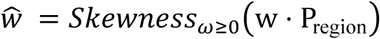

The weights were constrained to be non-negative to ensure meaningful scaling.

Fourthly, using the optimal weights *ŵ*, the final tumor purity was calculated as the mean of the weighted region purities:

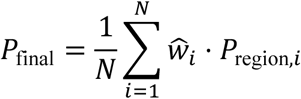

Here, *N* is the number of genomic regions, *ŵ_i_* is the optimal weight for the *i*-th region, and P_region,1_ is the initial purity estimate for that region.

### Attention score analysis in M2C tasks for significant CpG sites

To classify a sequence, the attention scores from all heads and all layers were collected and summed. Min-max scaling was then applied to obtain the scaled attention scores for all tokens. The attention scores were computed as follows:

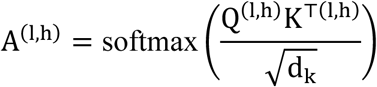

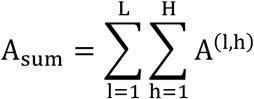

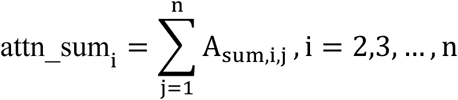

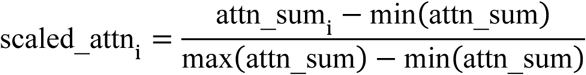

Here, *L* represents the number of encoder block layers, *H* denotes the number of attention heads per layer, *n* is the number of tokens of a sequence, and *d_k_* is the dimension of hidden state of an attention head. The [CLS] token was removed before applying min-max scaling.

For transcription factor (TF) binding motif identification at significant CpG sites, the reads located in the region of interest were collected and analyzed using HOMER (v5.1) with the findMotifsGenome.pl function and the parameter -size 150. If no associated motifs were identified, the 6 bp sequence surrounding the CpG site (in the format NNCGNN) was analyzed using Tomtom (v5.5.7, https://meme-suite.org/meme/tools/tomtom) to search for potential TF binding motifs. Sequences that were not associated with any HOMER- or Tomtom-enriched motifs, or motifs enriched without a sufficiently high CG bit score, were considered as possible novel motifs.

### Methylation to expression task fine-tuning

For the methylation-to-expression (M2E) task, ±2 kb regions flanking the transcription start site (TSS) of genes were considered as the input sequences. For TSS-flanking regions, CpG beta values were calculated using the formula:

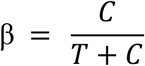

where *C* and *T* counts represented the sum of forward and reverse strand counts. Inside the sequence, Cs in CpGs with beta values ≥ 0.5 were converted to “M”, while those with beta values < 0.5 were converted to “C”. If a CpG site had no coverage, the corresponding sequence in the sample was discarded. The processed 4 kb methylation-encoded sequences were then paired with transcripts per million (TPM) values from matched RNA-seq libraries. Each pair of a methylation-encoded sequence and its corresponding TPM value was treated as a data point.

MethylQUEEN was fine-tuned using LoRA with mean squared error (MSE) loss. The fine-tuning process was performed with a batch size of 512, and a peak learning rate of 1e-5 with a linear warm-up of over 6% of the total training steps. Results were evaluated on the test dataset.

To supplement the training dataset, BS-seq libraries from the ENCODE and NIH Roadmap Epigenomics Project datasets were used. methylation-encoded sequence data from these samples were processed using the same method described above. The corresponding expression data were obtained from the GTEx database^28^. For each tissue, the mean and standard deviation of TPM values were calculated to construct a normal distribution of expression. For each methylation-encoded sequence, a paired TPM value was randomly sampled from this distribution. The fine-tuning process for the supplemented dataset followed the same procedure as described previously.

### Identification for potentially significant CpG sites for gene expression regulation

Potentially significant CpG sites were identified using attention scores when MethylQUEEN predicted expression values based on input methylation-encoded sequences. For each gene, all methylation-encoded sequences from the test set were analyzed. Scaled attention scores for all tokens were calculated using the same method described for the M2C task.

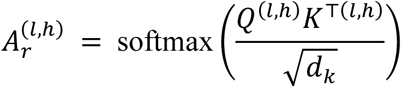

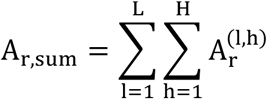

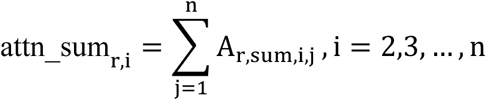

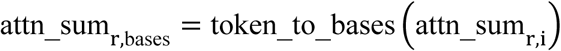

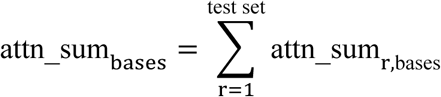

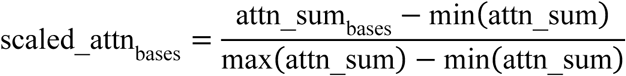

Here, *r* represent each piece of data in the test set. The function token_to_bases() converts the attention score of each token to each base of the sequence.

For all CpG sites, those with scaled attention scores ≥ 0.5 were classified as highly significant CpG sites, while those with scaled attention scores < 0.5 were classified as low-significant CpG sites.

## Data Availability

For methylation state prediction, the CSS HiFi PacBio dataset was constructed in this work, and will be available under publication.

For M2C task dataset, DLBCL vs B cell downstream task, Leucocyte deconvolution task, IMR-90 cell senescent vs Proliferiting task, and HCC vs healthy liver task data were from public datasets (GSE137880^38^, GSE186458^39^, GSE48580^40^, GSE70090^41^). For tumor burden estimation in plasma samples, target BS-seq libraries including HCC plasma samples from differenct stages and normal plasma from public dataset GSE149438^42^ were employed.

For the M2E task dataset, the matched RNA-seq and BS-seq dataset was from the public GEO database with accession GSE173790^43^.

## Code Availability

Upstream BS-seq data analysis pipeline is public on GitHub: https://github.com/Wubeizhongxinghua/DNA_5mC_analysis_pipeline.

The code and weights related to MethylQUEEN fine-tuning and inference will be made public after publication or under request.

## Acknowledgements

The authors would like to thank the support and computing resources provided by Polaris High Performance Computing platform (http://www.aais.pku.edu.cn/clshpc/) and computing platform in Peking University Chengdu Academy for Advanced Interdisciplinary Biotechnologies.

## Author Information

These authors contributed equally: Mingyang Li, Ruichu Gu.

## Contributions

Conceptualization: C.Y., M.L. and R.G. Supervision: C.Y., H.W., and B.H. Methodology, Investigation: M.L., R.G., S.F., Y.F., B.H., and J.Y. Software, Validation, Formal analysis, and Visualization: M.L., R.G., S.F, and Y.F. Resources: C.Y., H.W., B.H., M.L., Y.F., S.F., Y.C., M.X. Data Curation: M.L., S.F., Y.F. Writing - Original Draft: M.L., R.G. Writing - Review & Editing: C.Y., H.W., M.L., R.G., B.H., J.Y.

## Ethics declarations

The relevant patents are currently in the application process.

## Notes

### Summary of Updates

Competing Interests have been changed from “The authors have declared no competing interest.” to “The relevant patents are currently in the application process.”

